# Establishing a baseline for standardised genetic monitoring of Atlantic cod (*Gadus morhua*) in Sweden

**DOI:** 10.64898/2026.01.31.703008

**Authors:** Simon Henriksson, Carl André, Ricardo Tomás Pereyra, Håkan Wennhage, Kerstin Johannesson

## Abstract

Protecting populations and genetic diversity within them is critical to conserving the resilience and adaptive potential of species. Fisheries management has long had the ambition of managing species at the population level, but mainly define “fish stocks” based on geographical limits, which can lead to overfishing of sensitive populations in areas where many different populations coexist. Modern genetic methods are now sufficiently cost-effective, fast, and accurate to be integrated into fisheries management, enabling genetic identification and monitoring of fish populations. Here, we establish a genetic baseline for the commercially important fish Atlantic cod (*Gadus morhua*) in the waters surrounding Sweden, by using standardised sampling procedures and developing a genetic panel of 4000 single nucleotide polymorphisms (SNPs) for cost-effective assignment of population-of-origin and inversion genotypes. Using the SNP panel, we resolve the geographical distribution of three genetically distinct cod populations in the region: offshore, coastal/Western Baltic, and Eastern Baltic cod. While there is considerable spatial overlap between the three populations, they are genetically differentiated across the entire genome, as well as in genomic regions associated with chromosomal inversions. In addition, heterozygosity and effective population size estimates suggest differences in genetic diversity and rates of genetic erosion, underscoring the need to monitor the genetic diversity within each population separately. Repeating this methodology across years provides a first suggestion for establishing spatiotemporally resolved genetic monitoring of Atlantic cod in Sweden – simultaneously accounting for both population structure within the species and the genetic diversity within populations.

## 1. Introduction

Wild species are commonly subdivided into more or less discrete populations, and it is widely accepted that populations are the appropriate unit for conservation, rather than the species (Palsbøll *et al*., 2007; Reydon, 2019). While this has long been acknowledged in fisheries management (Hjort, 1914), management units, or “stocks”, are still mainly delineated geographically. If multiple fish populations coexist in an area, but are managed as a single stock, monitoring the overall stock size might not detect declines (or even local extinctions) of smaller populations unless population structure is accounted for (Cadrin, 2020). While it may not have been feasible before, genetic methods have now become accurate and inexpensive enough to employ within fisheries management (Andersson *et al*., 2024), enabling identification and monitoring of genetically differentiated fish populations (e.g., Johansen *et al*., 2018; Bekkevold *et al*., 2023).

Genetic diversity within species - both in the form of genetically distinct populations and genetic diversity within each population (Allendorf *et al*., 2022) - is a critical component of biodiversity, conferring resilience and adaptive potential to species in the light of short- and long-term environmental changes (Des Roches *et al*., 2021). In particular, genetic diversity associated with locally adapted populations, or “ecotypes”, is critical to maintain a species over a range of different environments (Johannesson *et al*., 2025). Genetic monitoring, where changes in population genetic parameters are assessed over time, can help identify losses of genetic diversity (Schwartz *et al*., 2007; Andersson *et al*., 2024), serving as an important indicator of the conservation status of populations and species (DeWoody *et al*., 2021). Genetic monitoring is, therefore, essential to safeguarding the persistence, resilience, and evolutionary potential of wild populations. Despite this, management programs that monitor changes in genetic diversity over time are very rare globally (Hunter *et al*., 2018), especially in marine environments (van Oppen & Coleman, 2022).

Atlantic cod (*Gadus morhua*, hereafter “cod”) is an ecologically and culturally important fish species, with multiple genetically differentiated populations and ecotypes described on both sides of the North Atlantic (e.g., Hemmer-Hansen *et al*., 2013; Kess *et al*., 2017; Knutsen *et al*., 2018). Cod is among the most well-studied species in the Northeast Atlantic with regard to connectivity and population structure (Henriksson *et al*., 2025), and multiple genetic SNP panels for population assignment have been developed (e.g. Kent *et al*., *in prep*.; Weist *et al*., 2019; Hemmer-Hansen *et al*., 2019). Together, these factors make cod a highly suitable species to establish genetic monitoring of. Such genetic monitoring has already been implemented for cod populations in other regions, informing local fisheries management (e.g., Dahle *et al*., 2018; Christensen *et al*., 2022).

In Sweden, three genetically distinct cod populations are currently described (Figure 1): “offshore” (or “North Sea”) cod, “coastal” (or “Kattegat” or “Western Baltic”; WBC) cod, and “Eastern Baltic” cod (EBC). The nomenclature used for each genetic population differs across studies, depending on their geographic scope. On the Swedish west coast, the two former populations are often referred to as the “offshore” and “coastal” ecotypes, as they are in part associated with different environments: deep offshore and shallow coastal habitats, respectively (Henriksson *et al*., 2023). Both ecotypes coexist on small spatial scales (Knutsen *et al*., 2018) and both can spawn inside the same fjords (Jorde *et al*., 2018; Svedäng *et al*., 2019). Despite spawning both offshore and inside Skagerrak fjords, the offshore ecotype cod found in the fjords has not been genetically distinguished from cod spawning in the North Sea (Barth *et al*., 2019). Similarly, the coastal ecotype cod found in Swedish fjords does not appear genetically distinct from cod in the Kattegat or Western Baltic Sea (Barth *et al*., 2017; Henriksson *et al*., 2023). While genetically distinct “fjord” cod is documented in Norwegian Skagerrak fjords (Barth *et al*., 2019), cod is strongly depleted in Swedish fjords (Svedäng 2003), and it is still unclear whether “fjord” cod, distinct from the offshore and coastal ecotypes, exists in Sweden (*cf*. Henriksson *et al*., 2023). In the Baltic Sea, WBC and EBC coexist in the Arkona basin (ICES Subdivision 24), but are otherwise separated geographically and have non-overlapping spawning periods (Hemmer-Hansen *et al*., 2019). Analogous to the ecotypes on the Swedish west coast, WBC occupies shallower depths than EBC in areas where they overlap (Schade *et al*., 2022). Based on the geographic area of the current study, we hereafter refer to the three populations as “offshore”, “coastal/WBC” and “EBC” cod.

**Figure 1.**
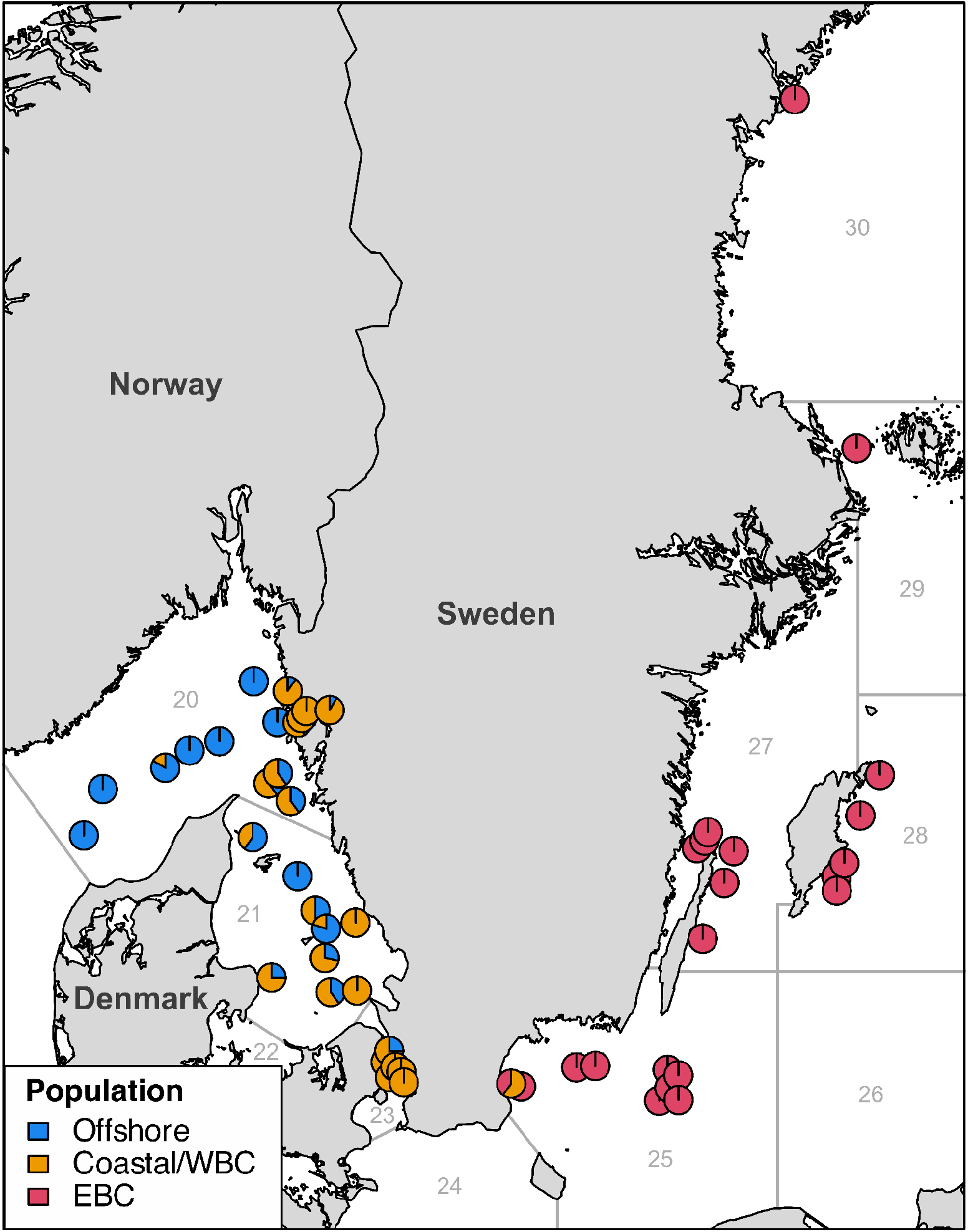
The geographic distribution of individuals from the three populations in Swedish waters: offshore cod, coastal/Western Baltic cod (WBC), and Eastern Baltic cod (EBC). Each pie chart indicates the relative proportions of each population at each locality. Numbers denote ICES subdivisions.

Across the Skagerrak-Kattegat-Baltic Sea region, the relative proportions of different cod populations appear highly variable across different years (Synnes *et al*., 2021; Schade *et al*., 2022), within years (Jorde *et al*., 2018), and across age groups (Hemmer-Hansen *et al*., 2020). This can be attributed to variability in the source of recruits across years (Cardinale & Svedäng, 2004; Knutsen *et al*., 2018; Jonsson *et al*., 2016), as well as complex migratory behaviours of adults (André *et al*., 2016; Mion *et al*., 2022; Lundgreen *et al*., 2023). Standardised and temporally replicated genetic monitoring would further develop our understanding of the complex spatiotemporal dynamics of Swedish cod populations, fundamental to establishing population-based management practices in the region.

A major challenge to establishing genetic monitoring programs lies in the standardisation of sampling and genotyping methods across years. Sampling of wild fish species can be labour-intensive and expensive, making it difficult to design sampling methodologies that are simultaneously both representative, standardised, and cost-effective. Such sampling efforts are, however, already in place within the national fisheries-independent surveys aimed at quantitative monitoring of fish stocks (e.g., Bland & Börjesson, 2020; Lövgren, 2020). Furthermore, the rapid development of molecular methods means that new and improved genetic tools are constantly being introduced. Although this progress has led to the development of tools that can now accurately distinguish fish populations (Andersson *et al*., 2024), it simultaneously makes it challenging to standardise which genetic methods to apply within genetic monitoring programs, which need to be consistent across years. As the amount of genetic information on wild populations increases, however, researchers have a responsibility to make genetic resources (data, information, and tools) readily available for practical implementation in management (Hunter *et al*., 2018; Hoban *et al*., 2020).

Here, we describe a genetic SNP panel designed for population genetic analysis of Atlantic cod in the waters surrounding Sweden. We demonstrate its utility in both establishing a genetic baseline for monitoring of cod populations in Sweden and in assigning individual cod in smaller research projects to their population of origin through comparison to this baseline.

## 2. Methods

### 2.1. Sample collection

In total, tissue samples from 474 individual cod were collected (Table 1) by the Swedish University of Agricultural Sciences (SLU). A large fraction of these cod (n=210) was collected during fisheries-independent surveys in 2020 as part of the annual quantitative monitoring of fish stocks. These include samples from offshore and coastal Skagerrak (SKG_OFF, n=30; SKG_COAST, n=30) and offshore Kattegat (KTG_OFF, n=40) collected in the International Bottom Trawling Survey (IBTS), samples from Öresund (ORE, n=30) collected during the Swedish Sound Survey, as well as samples from the Hanö Bay (HAN, n=20), southern Midsjöbank (BANK, n=20), Öland (OLAND, n=20), and Gotland (GOT, n=20) collected in the Baltic International Trawl Survey (BITS). To extend the geographic coverage in the northern Baltic Sea, we also included samples from Åland (AL, n=100) collected in 2019 (n=40) and 2021 (n=60), and from the Bothnian Sea collected in 2016 (BH, n=19). To increase the geographic coverage in inshore Skagerrak, which currently is not subject to regular fisheries-independent surveys, we included samples from Bovallstrand (BVS, n=23), inner Gullmarsfjorden (GUL, n=7), outer Gullmarsfjorden (CLIM, n=49, used in Perry *et al*., 2024), and Byfjorden (BF, n=66). Detailed metadata is provided in Table S1.

**Table 1.** List of samples included in the study.

| Location | Code | ICES<br>Subdivision | Years | N |
| --- | --- | --- | --- | --- |
| Skagerrak offshore | SKG_OFF | 20 | 2020 | 30 |
| Skagerrak coast | SKG_COAST | 20 | 2020 | 30 |
| Bovallstrand | BVS | 20 | 2013-2021 | 23 |
| Outer Gullmarsfjorden | CLIM | 20 | 2020 | 49 |
| Inner Gullmarsfjorden | GUL | 20 | 2013-2014 | 7 |
| Byfjorden | BF | 20 | 2019-2021 | 66 |
| Kattegat offshore | KTG_OFF | 21 | 2020 | 40 |
| Öresund | ORE | 23 | 2020 | 30 |
| Hanö bay | HAN | 25 | 2020 | 20 |
| Södra Midsjöbanken | BANK | 25 | 2020 | 20 |
| Öland | OLAND | 27 | 2020 | 20 |
| Gotland | GOT | 28 | 2020 | 20 |
| Åland | AL | 29 | 2019-2021 | 100 |
| Bothnian Sea | BH | 30 | 2016 | 19 |

### 2.2. Genotyping

DNA extraction and genotyping of all 474 cod was performed by IdentiGEN LTD., Dublin, Ireland. For genotyping, we used a new SNP chip designed for diagnostic population assignment in multiple fish species – Atlantic herring (*Clupea harengus*), horse mackerel (*Trachurus trachurus*), Atlantic salmon (*Salmo salar*), brown trout (*Salmo trutta*), European perch (*Perca fluviatilis*), and cod (MultiFishSNPChip_1.0; Andersson *et al*., 2024). The chip includes 3000-4000 SNP loci per species, and has already been applied in studies on perch (Vasemägi *et al*., 2023), herring (Pettersson *et al*., 2024; Goodall *et al*., 2024), and cod (Perry *et al*., 2024). Below we describe how we designed the SNP panel for population assignment in cod.

A detailed list of the SNP loci used in the cod SNP panel is given in Table S2. The majority of loci originate from a panel of 10 605 SNPs (Kent *et al*., *in prep*.), used in multiple population genetic studies on cod in the North Atlantic (e.g., Berg *et al*., 2015; Sodeland *et al*., 2016; Barth *et al*., 2017). We also included loci from an ecotype- and inversion-diagnostic panel used for cod ecotypes on the Swedish west coast (Henriksson *et al*., 2023), as well as previously identified global outlier loci for Baltic cod (Berg *et al*., 2015), which enable investigation of potential substructure in eastern Baltic cod. To further increase the discriminatory power of our genetic panel, we added loci from two separate SNP panels designed for distinguishing WBC and EBC (Hemmer-Hansen *et al*., 2019; Weist *et al*., 2019).

All loci (n=10 674) were re-mapped to the gadMor3.0 reference genome (Jentoft *et al*., 2025) based on sequence information on the flanking regions presented in the original publications. Mapping was performed using BowTie2 (v2.3.4.3; Langmead & Salzberg, 2012), and loci that were mapped more than once, or were duplicated across genetic panels, were removed. After additional thinning based on alignment quality and *in vitro* amplification of the markers, the final panel consisted of 4032 loci – 3786 from Kent *et al*. (*in prep*.), 152 from Berg *et al*. (2015), 29 (including 9 inversion-diagnostic loci) from Henriksson *et al*. (2023), 21 from Weist *et al*. (2019), 12 from Hemmer-Hansen *et al*. (2019), and 32 control loci. The loci from Berg *et al*. (2015), Hemmer-Hansen *et al*. (2019), Weist *et al*. (2019), and Henriksson *et al*. (2023) were repeated 8x on the chip, to maximise the chances of successful genotyping at these highly discriminatory loci. To ensure that population clustering followed expectations and was consistent with previous assignments, 27 individuals with known genetic origins from Henriksson *et al*. (2023) were included among the 474 individuals in this study.

### 2.3. Genetic analysis

#### 2.3.1. Quality control and filtering

Individuals and loci with call rates below 90% were removed from downstream analysis. We also filtered out potentially contaminated samples, identified by calculating mean kinship and heterozygosity for all individuals, using the SNPRelate (v1.38.0; Zheng *et al*., 2012) and dartR packages (v2.9.7; Gruber *et al*., 2018), respectively. Individuals that had both mean kinship and heterozygosity values in the 95^th^ percentile and clustered together in the kinship analysis were removed.

#### 2.3.2. Inversion scan

The Atlantic cod genome contains several chromosomal inversions (Matschiner *et al*., 2022; Hoff *et al*., 2024). Loci within these genomic regions are under high linkage disequilibrium (LD), which masks the underlying population structure unless it is accounted for. To identify the genomic regions where chromosomal inversions are located, we used the inveRsion package (v1.40.0, Cáceres, 2021) to perform scans across each chromosome (haplotype block size: 3 SNPs; window size: 1-30 Mb; iterations: ≤30; MAF: 5%; BIC: >0). To avoid incompatibility issues across R packages, this analysis was run in R v4.1.2 (R Core Team, 2021). This method successfully detected the three inversions on chromosomes 2, 7, and 12 that are polymorphic in this geographic region (e.g., Sodeland *et al*., 2022). As the 4000 SNP loci provide a somewhat limited coverage for genomic scans such as these, we compared the breakpoints inferred herein against breakpoints inferred from a dataset with genotypes at 9 956 loci (Henriksson *et al*., 2023). This comparative approach demonstrated a high degree of consistency between datasets, and the breakpoints flanked genomic blocks of loci with high LD (Figure S1), calculated with the dartR package (Gruber *et al*., 2018). To delineate the inverted regions in downstream analyses, we have conservatively used the most extreme break points out of these options, to minimise the risk of including any loci linked to the inversions in other downstream analyses. As the inversion on chromosome 1 was not polymorphic in our populations, we used breakpoints previously inferred from whole-genome data (Aasegg Araya *et al*., 2025). The inversion scan also identified a smaller inversion on chromosome 4, which has been described in previous studies (Barth *et al*., 2017; Hoff *et al*., 2024), and showed clear trimodal clustering in a PCA, consistent with that of a chromosomal inversion (Ma & Amos, 2012). Because of this, this chromosomal region was also treated as an inversion in downstream analyses. We identified additional small regions with potential inversions on all chromosomes apart from chromosomes 5, 7, 14, 20, and 22 (see Table S3; Figure S1). Although of interest for future studies on chromosomal inversions, these smaller inversions will not be discussed in detail here. Loci within the inverted regions on chromosomes 1, 2, 4, 7, and 12 were removed in all downstream analyses that did not explicitly analyse inversion genotypes, to minimise issues with LD. We will refer to loci outside of these inverted regions as “collinear” loci.

#### 2.3.3. Assignment of population-of-origin and inversion genotypes

Assignment of individuals to population was performed *de novo* using the K-means clustering algorithm in the adegenet package (v2.1.11; Jombart, 2008). The resulting assignments were compared for 27 individuals assigned in a previous study (Henriksson *et al*., 2023), with consistent population assignment for 100% of individuals (Figure S2A). For population assignment, we used the full panel of collinear loci, as this panel gave the clearest clustering in principal component analyses (PCA) and the highest posterior probabilities for the assignments in a discriminant analysis of principal components (DAPC; adegenet; Jombart 2008). To investigate potential hybridisation among populations, we calculated ancestry coefficients using sparse non-negative matrix factorisation (sNMF) in the LEA package (v3.16.0; Frichot & François, 2015).

The assignment of individual inversion genotypes was performed by PCA with the adegenet package, based on loci located inside each inverted region on chromosomes 2, 4, 7, and 12. Individual inversion genotypes showed clear trimodal clustering along PC1, and were assigned following Ma & Amos (2012). Inversion genotypes for chromosomes 2, 7, and 12 were assigned using the nomenclature of Matschiner *et al*. (2022; *cf*. Henriksson *et al*., 2023). For the inversion on chromosome 4 that was not included in this nomenclature, we have named the most frequent arrangement as “Homozygote 1”, and the alternate arrangement as “Homozygote 2”.

#### 2.3.4. Genetic divergence and diversity

To assess genetic divergence between populations, we estimated pairwise *F*_ST_ (Weir & Cockerham, 1984) at each individual SNP locus, as well as for all collinear loci (10 000 permutations), using the strataG package (v2.5.1; Archer *et al*., 2016). We furthermore estimated the effective population sizes (*N*_e_) of each population based on LD, using the strataG package. To estimate *F*_ST_ among populations for each inversion, the inferred genotypes were re-coded as single superlocus genotypes before calculating *F*_ST_ as described above. To infer potential adaptive divergence, we performed outlier analysis for all population pairs using OutFLANK with default settings (v0.2; Whitlock and Lotterhos, 2014). Observed heterozygosity (*H*_o_) was calculated using the hierfstat package (v0.5-11; Goudet & Jombart, 2022). We used *H*_o_ instead of *H*_e_ as this enabled estimating heterozygosity across multiple loci replicated per individual, thus, providing a range of heterozygosity within each population. Statistical differences in *H*_o_ between populations were tested using analysis of variance (ANOVA) with the car package (v.3.1.2; Fox & Weisberg, 2019), followed by a Tukey Honest Significant Differences (HSD) post-hoc test, using the stats package.

Unless otherwise stated, all population genetic and statistical analyses were performed with R (v4.4.0, R Core Team, 2024) in RStudio (v2024.4.1.748; Posit team, 2024).

## 3. Results and discussion

### 3.1. Genotyping success

The newly developed SNP panel produced genotypes for 457 out of 474 individuals (96.4%) with a call rate >90% for 3776 out of 4000 loci (94.4%). A further nine individuals were removed due to suspected contamination (high *H*_o_ and high mean kinship). The kinship analysis also identified two genetically identical individuals from inner Gullmarsfjorden, most likely due to fin-clipping the same individual twice during field sampling (in 2013-2014). This example demonstrates the applicability of this panel also in wildlife forensics and individual recognition. One of these individuals was removed from downstream analysis.

### 3.2. Genetic baseline

#### 3.2.1. Population structure

The genetic panel clearly identifies the three populations previously documented in the waters around Sweden. Overall, our samples included 86 offshore cod, 172 coastal/WBC cod, and 189 EBC. The populations cluster into discrete groups in the PCA (Figure 2), supported by high posterior assignment probabilities in the DAPC. As described in previous studies, offshore cod was primarily associated with offshore sites, whereas the coastal/WBC cod was the most frequent in coastal and inshore Skagerrak, as well as in the Kattegat and Öresund (Figure 1). The high proportion of coastal/WBC cod in the offshore regions of eastern Skagerrak, however, contrasts slightly with previous population genetic studies (Henriksson *et al*., 2023). This discrepancy points to variability in the relative abundance of the two populations across years and age groups, as has been shown in the Kattegat (Hemmer-Hansen *et al*., 2020), and further underscores the need for temporally resolved genetic monitoring also in offshore Skagerrak. Meanwhile, the Baltic Proper is almost exclusively occupied by EBC, while three individuals of coastal/WBC cod were also found in the Hanö bay, north of the Danish island Bornholm. This sample was taken close to the border of ICES Subdivision 24, where both WBC and EBC occur in sympatry (e.g., Schade *et al*., 2022). Sweden does not regularly perform fisheries-independent surveys of fish stocks in ICES Subdivision 24, and, hence, no samples from there were included in this study. No EBC were found outside of the Eastern Baltic Sea, consistent with tagging studies showing limited movement of EBC outside of the Baltic Sea (e.g., Mion *et al*., 2022). Similarly, the distribution of coastal/WBC cod across subdivisions 21-25 is supported by movement patterns of cod captured and released in Öresund (Western Baltic Sea; Lundgreen *et al*., 2023).

**Figure 2.**
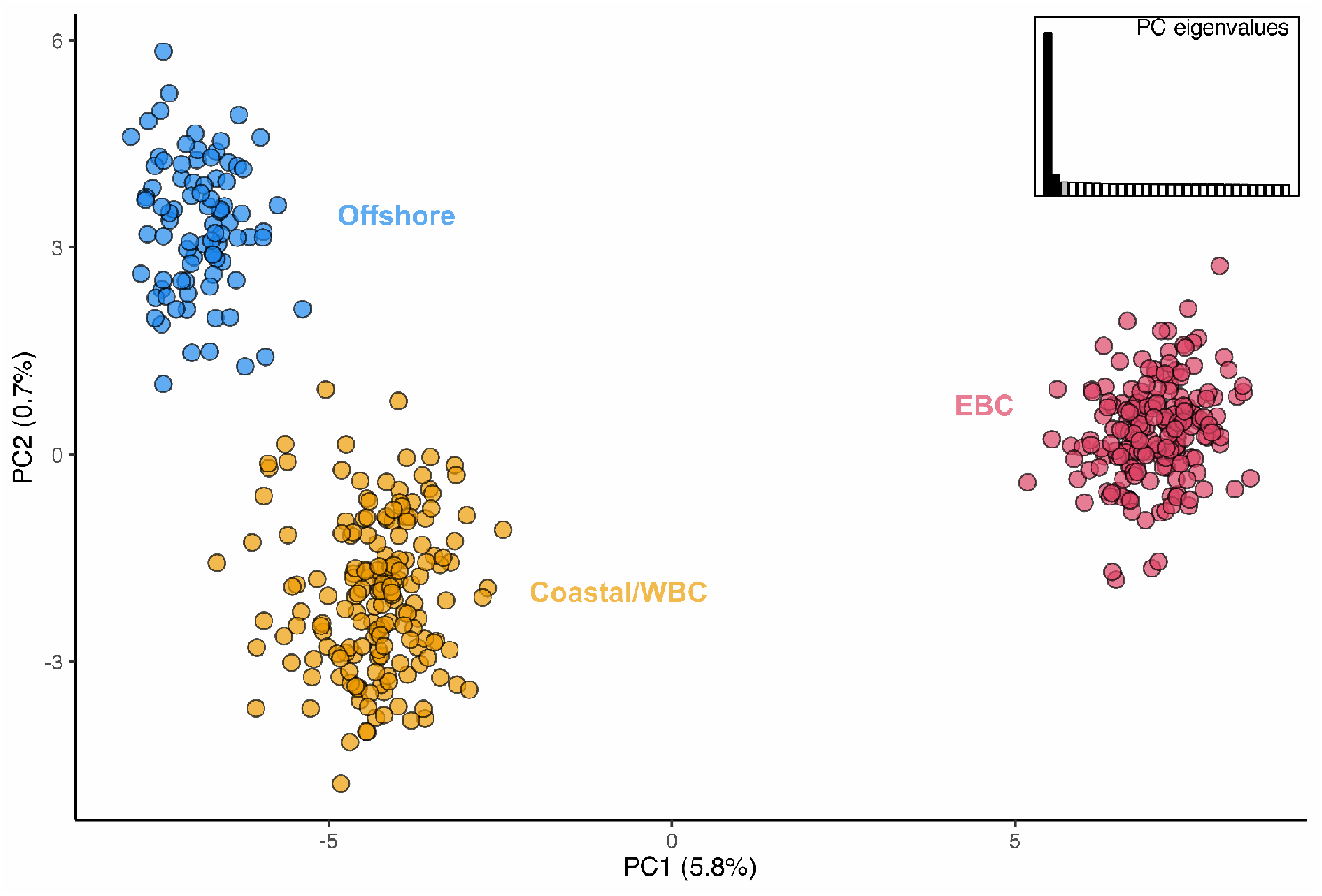
Principal component analysis (PCA) based on genetic variation at collinear loci. Colour corresponds to the population of each individual, as assigned through *K*-means clustering - offshore cod (blue), coastal/Western Baltic cod (WBC; orange), and Eastern Baltic cod (EBC; pink). The scree plot displays eigenvalues of PCs 1-30, and the percentage of variation explained by the first and second PC is given within parenthesis on the x and y axis, respectively.

#### 3.2.2. Genomic divergence

Both the clustering and ancestry analyses suggest very limited gene flow between EBC and the two western populations. This supports previous evidence for EBC being highly distinct from adjacent populations (e.g., Berg *et al*., 2015; Barth *et al*., 2019). This was also supported by genetic divergence (*F*_ST_) between EBC and both western populations being higher than divergence between the offshore and coastal/WBC populations, at collinear loci (Table 2). The ancestry analysis indicated that some genetic admixture may occur between offshore and coastal/WBC cod (Figure S3), however, first-generation hybrids appear very rare. Locus-wise *F*_ST_ between offshore and coastal/WBC cod was highest at loci within the chromosomal inversion on chromosome 12 (Figure 3A), while being high across the whole genome for EBC compared to the other two populations (Figure 3B-C). The outlier analyses support the more heterogeneous genomic divergence pattern of offshore and coastal/WBC cod, with 157 outlier loci detected, out of which 87 (55%) were located within the inversion on chromosome 12. The comparison of EBC to the offshore and coastal/WBC cod returned fewer outlier loci (coastal/WBC vs EBC: n = 80; offshore vs EBC: n = 11), which were more widespread across the genome. The smaller number of outlier loci is probably due to EBC having high overall *F*_ST_ against the other two populations. The difference in the number of outlier loci inferred in each pairwise comparison is, in part, also attributable to the differences in sample size among populations (see Section 3.2.1). Note that the genetic panel consists of SNP loci known to be divergent across these three populations, and the *F*_ST_ estimates presented herein would likely differ from those of a random set of loci.

**Table 2.** Pairwise genetic divergence *F*_ST_ (Weir & Cockerham, 1984) between populations (OFF = offshore; COA = coastal/Western Baltic; and EBC = Eastern Baltic cod) for collinear loci, and for each inversion genotype. Uncorrected and experiment-wise corrected p-values (q-vsalues) are shown, and comparisons with significant q-values are shown in bold font.

| | Comparison | $F_{ST}$ | p-value | q-value |
| --- | --- | --- | --- | --- |
| Collinear loci | OFF vs COA | 0.011 | <0.001 | 0.002 |
|  | COA vs EBC | 0.052 | <0.001 | 0.002 |
|  | OFF vs EBC | 0.078 | <0.001 | 0.002 |
| Chr 2 inversion | OFF vs COA | 0.069 | <0.001 | 0.002 |
|  | COA vs EBC | 0.487 | <0.001 | 0.002 |
|  | OFF vs EBC | 0.613 | <0.001 | 0.002 |
| Chr 4 inversion | OFF vs COA | 0.031 | 0.008 | 0.041 |
|  | COA vs EBC | 0.468 | <0.001 | 0.002 |
|  | OFF vs EBC | 0.575 | <0.001 | 0.002 |
| Chr 7 inversion | OFF vs COA | 0.008 | 0.104 | 0.313 |
|  | COA vs EBC | 0.004 | 0.137 | 0.313 |
|  | OFF vs EBC | -0.003 | 0.691 | 0.691 |
| Chr 12 inversion | OFF vs COA | 0.183 | <0.001 | 0.002 |
|  | COA vs EBC | 0.082 | <0.001 | 0.002 |
|  | OFF vs EBC | 0.017 | 0.031 | 0.123 |

**Figure 3.**
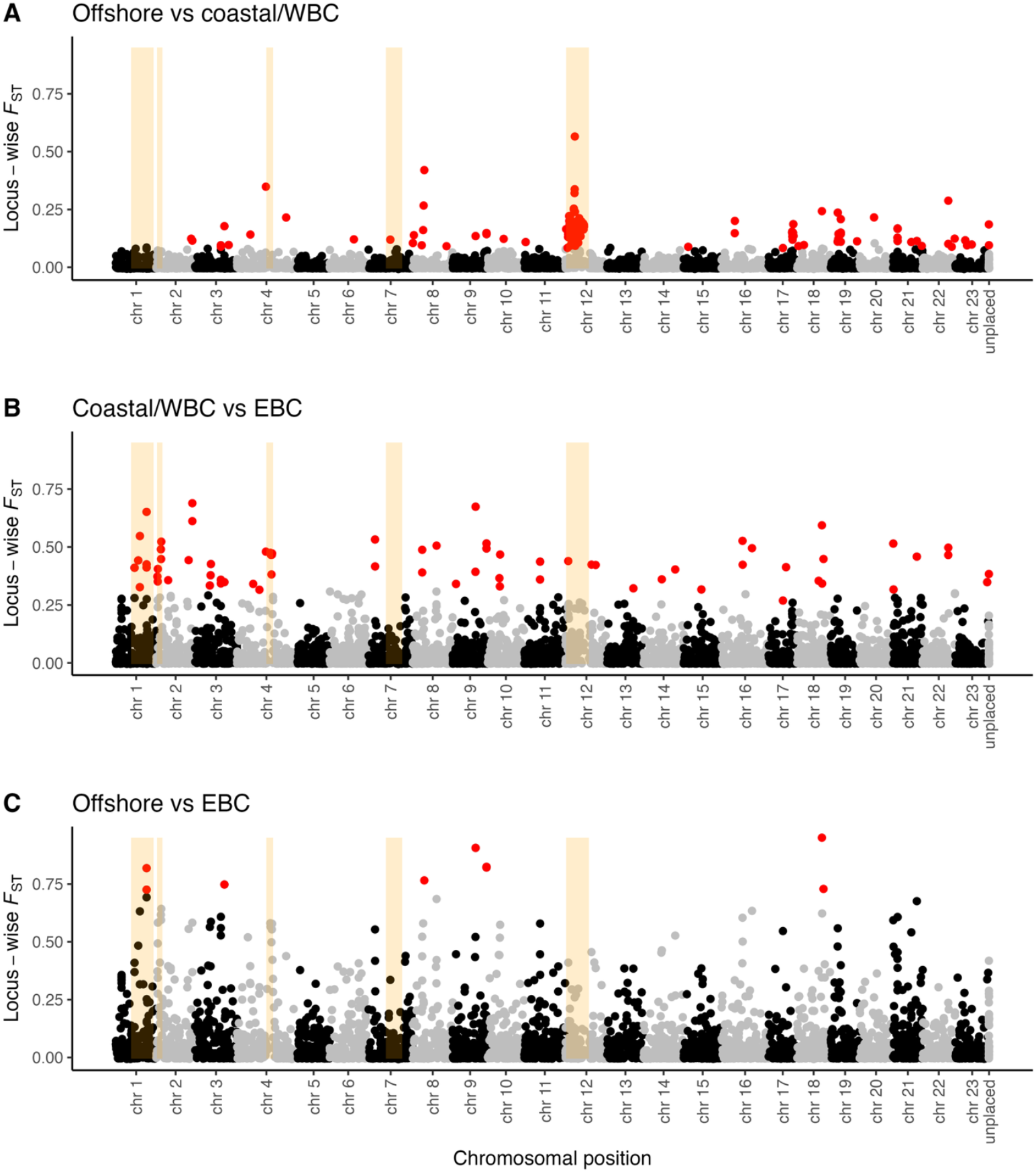
Pairwise *F*_ST_ per locus between the three cod populations: **A)** offshore and coastal/WBC, **B)** coastal/WBC and EBC, and **C)** offshore and EBC. The inverted regions on chromosomes 1, 2, 4, 7, and 12 are highlighted in orange, and statistical outlier loci are indicated with red points.

Inversion genotype frequencies differed significantly among populations (Table 2; Figure 4). The inversions on chromosomes 2 and 4 had very similar divergence patterns among populations, and *F*_ST_ values were significantly different for all pairwise comparisons. The divergence at the chromosome 2 and 4 inversions was roughly one order of magnitude higher for EBC compared to offshore or coastal/WBC cod, than between the offshore and coastal/WBC cod. For the inversions on chromosomes 2 and 4, however, there were also consistent gradients in genotype frequency from the south to the north in the Eastern Baltic Sea (Figure 5; S4), suggesting adaptive divergence within EBC, potentially linked to salinity adaptation (Berg *et al*., 2015). In contrast, the chromosome 7 inversion did not differ across populations. The chromosome 12 inversion also did not differ between offshore cod and EBC, but did so for both these populations compared against coastal/WBC cod. Interestingly, however, the difference in *F*_ST_ was twice as large between the highly sympatric offshore and coastal/WBC cod as between coastal/WBC and EBC cod. Cod homozygous for the derived arrangement on chromosome 12 have previously been shown to have higher survival in Norwegian fjords (Barth *et al*., 2019). The divergence of coastal/WBC cod against both offshore cod and EBC at this inversion, thus, suggests a functional role of genes located on chromosome 12 in adaptation to coastal environments.

**Figure 4.**
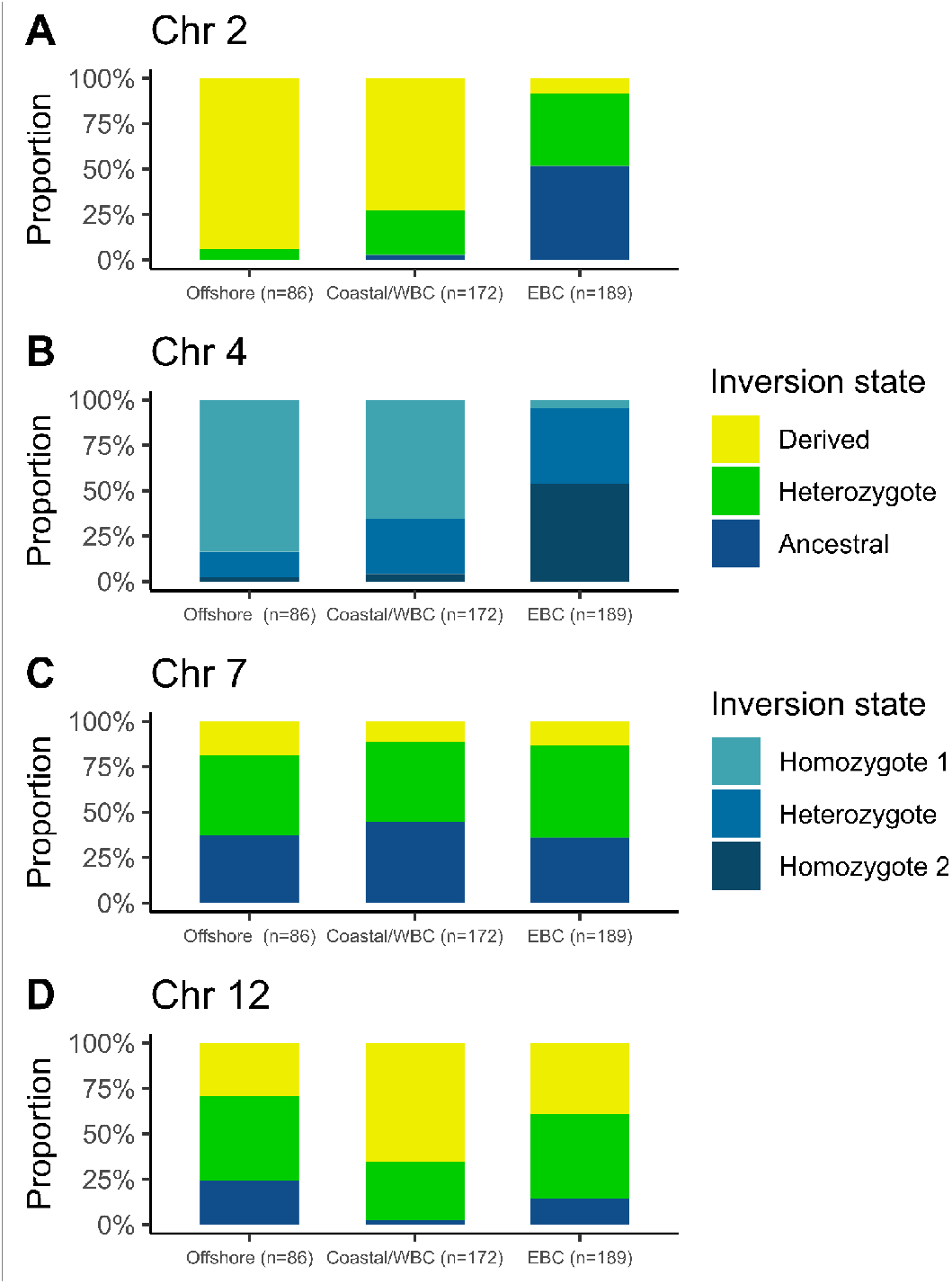
Inversion genotype frequencies for inversions on chromosomes **A)** 2, **B)** 4, **C)** 7, and **D)** 12 per population. Note that the ancestral and derived arrangements are unknown for the inversion on chromosome 4.

**Figure 5.**
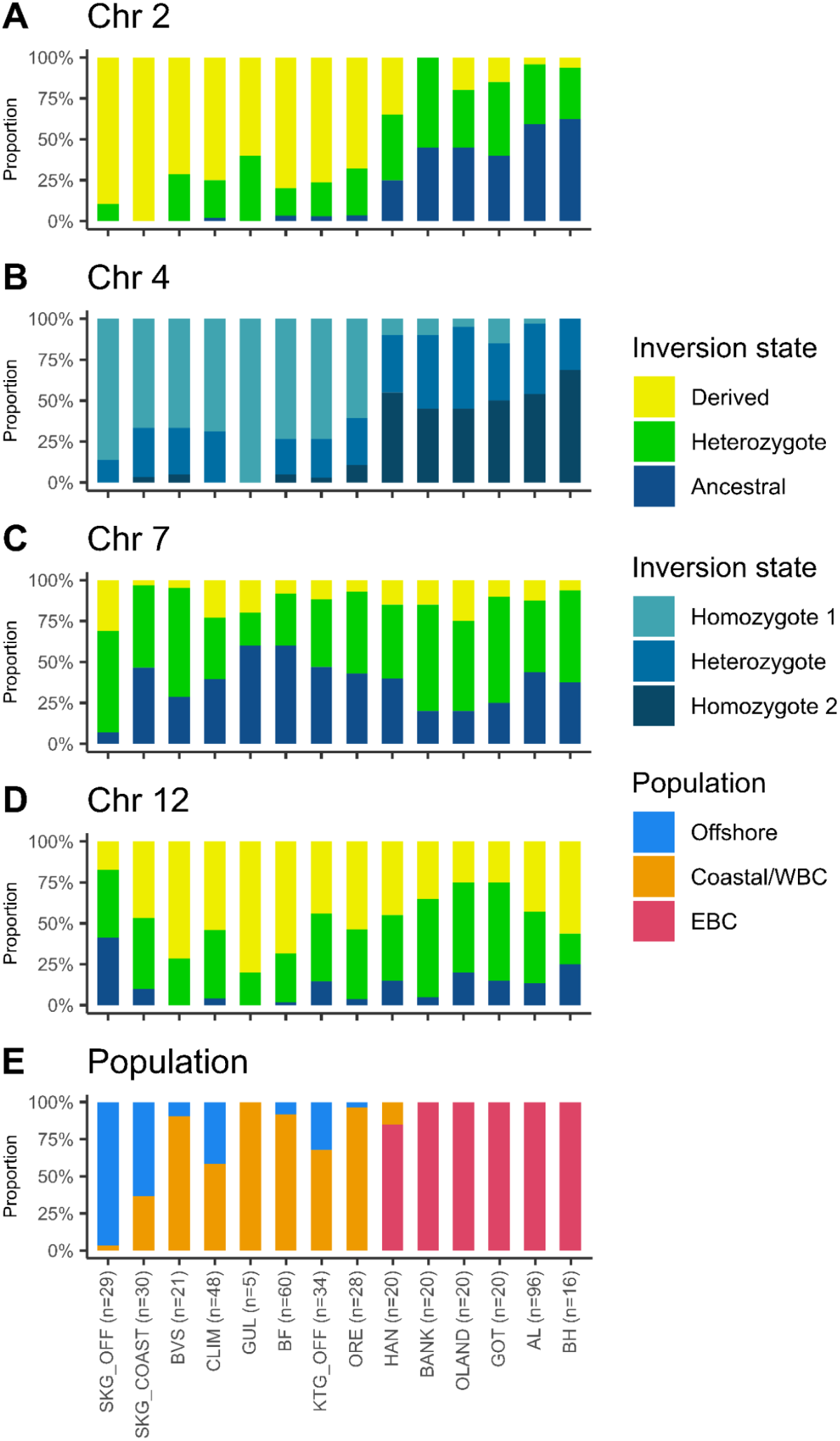
Genotype frequencies of the chromosomal inversions on chromosomes **A)** 2, **B)** 4, **C)** 7, and **D)** 12, as well as **E)** population proportions, at each location. All locations are sorted from offshore Skagerrak (left), southwards into the Baltic, turning northwards for the Baltic samples (right).

Lastly, offshore and coastal/WBC cod showed a distinct peak in *F*_ST_ in the central region of the chromosome 12 inversion (position ∼6.645-7.066 Mb in GadMor3.0; Figure S5A). This region has been subject to a double crossover event in EBC and harbours genes thought to be involved in reproduction and egg buoyancy regulation (Matschiner *et al*., 2022). The same peak in genomic divergence between offshore and coastal/WBC cod has been observed previously (Henriksson *et al*., 2023), but was not pronounced in either of the pairwise comparisons against EBC in the current study (Figures S5B-C).

Overall, the results show genome-wide divergence among the three populations, but also more pronounced divergence in certain genomic regions potentially linked to environmental adaptation and ecotype formation. The Skagerrak-Baltic Sea region is characterised by multiple environmental gradients at both large and small spatial scales (Snoeijs-Leijonmalm & Andrén 2017). Managing Atlantic cod in accordance with this population structure might, therefore, be of critical importance for the persistence of the species in Sweden.

#### 3.2.3. Genetic diversity within populations

Our estimates of *H*_o_ and *N*_e_ within each population gives insight to the genetic diversity and evolutionary histories of the three populations (Figure 6). By replicating the current study temporally, using the same genetic panel, changes in genetic diversity can be monitored over time. All populations had significantly different *H*_o_ values (q < 0.01; Table S4), with the highest values in offshore cod (mean *H*_o_ = 0.350), followed by coastal/WBC cod (mean *H*_o_ = 0.347) and EBC (mean *H*_o_ = 0.317). The pattern was reflected in the *N*_e_ estimates (offshore: mean *N*_e_ = 7916, CI = 5206-16457; coastal: mean *N*_e_ = 6017, CI = 4740-8226; EBC: mean *N*_e_ = 5261, CI = 4213-6991), although differences in *N*_e_ were non-significant. The results suggest gradually decreasing genetic diversity going from offshore Skagerrak into the Baltic Sea. Lower heterozygosity in EBC is consistent with a founder effect, which would align with the proposed evolutionary history of EBC as having diverged from coastal/WBC cod, perhaps as recently as ∼10 000 years ago (Matschiner *et al*., 2022). Simultaneously, if EBC has lower *N*_e_ than the other populations, this population will lose genetic diversity more rapidly due to genetic drift, although still at a slow rate at the current size of *N*_e_. This result highlights the need to monitor the cod populations separately, as genetic erosion within each population may occur at different rates. Because of its distinct environmental adaptation and reproductive isolation from adjacent populations, EBC was originally described as its own subspecies of cod (Linnaeus, 1761), however, hybridisation between WBC and EBC has occurred in periods when the Baltic cod population sizes were large (Helmerson *et al*., 2023).

**Figure 6.**
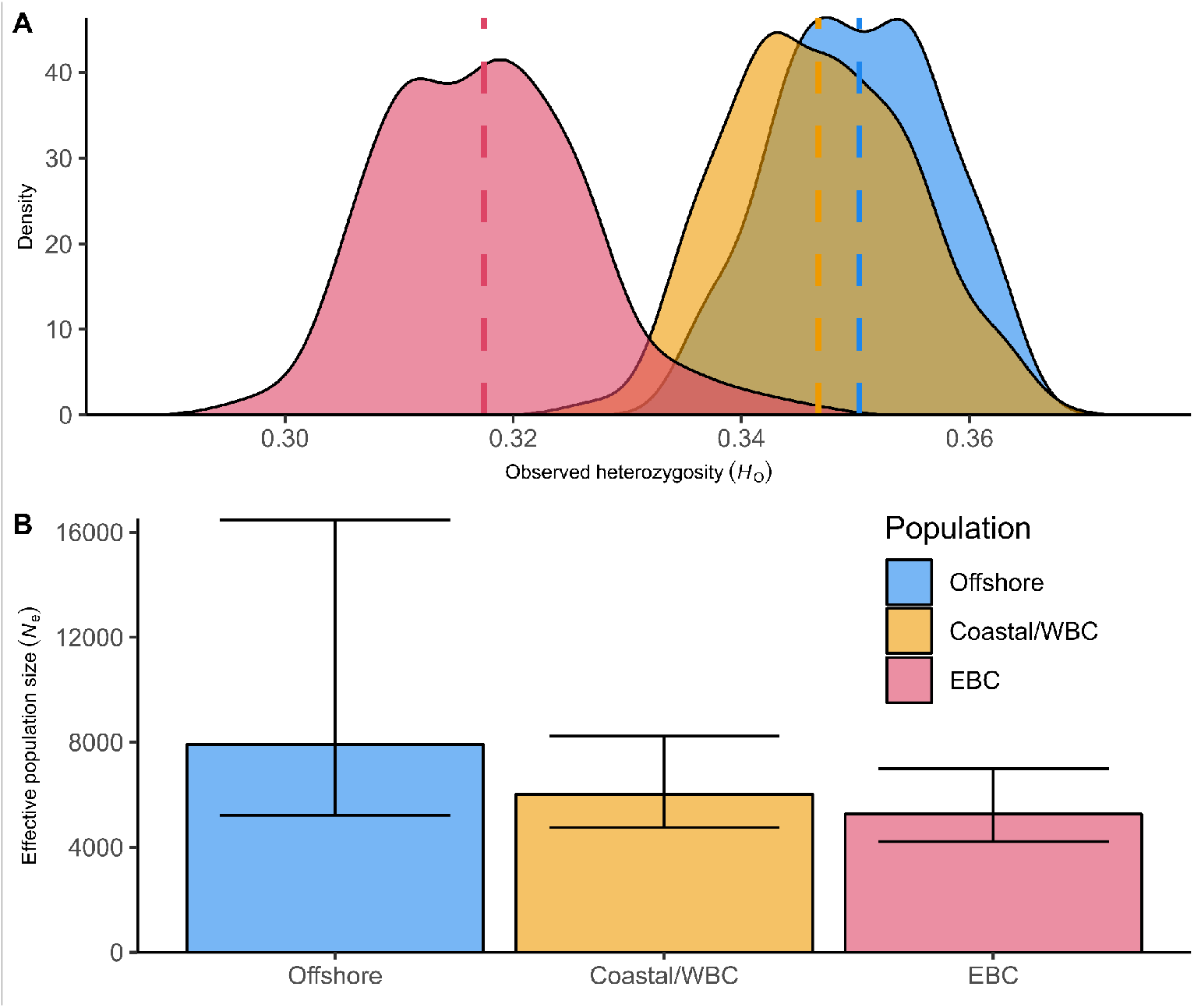
Genetic diversity within populations. **A)** Observed heterozygosity (*H*_o_) and **B)** effective population size of each population. Dashed lines in **A** indicate the mean *H*_o_ for each population, and error bars in **B** show the 95% confidence intervals for the *N*_e_ estimates. All estimates exclude loci within inversions.

### 3.3. Case studies

Some of the samples included in the genetic baseline dataset presented herein originate from scientific studies with different aims and smaller numbers of samples. These include cod from an experiment investigating how cod responds to climate stressors, and a telemetry study investigating the movement ecology of large cod in inshore Skagerrak. The samples collected from Åland and the Gulf of Bothnia were also of special interest for investigations of potential population substructure within EBC. Below we provide an overview of a few smaller “case studies”, which demonstrate the utility of using this SNP panel and genetic baseline in future studies.

#### 3.3.1. Climate stressor experiment

The 49 wild-caught cod included in the climate stressor experiment were collected in outer Gullmarsfjorden in 2020, and 48 were successfully genotyped. Out of these, 20 (42%) were of the offshore ecotype, and 28 (58%) were of the coastal ecotype (Figure S2B). The physiological results from this experiment are presented in detail by Perry *et al*. (2024), but, in brief, the authors showed that coastal ecotype cod were more resistant to climate stressors (high temperatures and low salinities) than offshore cod were. Genotypes for these fish are previously deposited on the Zenodo data repository (https://doi.org/10.5281/zenodo.10075426).

#### 3.3.2. Large cod in inshore Skagerrak

Among the 66 larger cod captured in Byfjorden, 60 were successfully genotyped. Five (8%) were of the offshore ecotype, and the remaining 55 (92%) were of the coastal ecotype (Figure S2C). This contrasts with a previous study on juvenile cod sampled in 2019 and 2020, estimating that 17-44% of juveniles are of offshore origin in Byfjorden (Henriksson *et al*., 2023). A similarly high proportion of coastal cod (90%) was observed among the larger cod in Bovallstrand (BVS; n=21 successfully genotyped cod). These results support the notion that some proportion of offshore cod leave coastal Skagerrak and Skagerrak fjords upon reaching a certain size or age, whereas the coastal cod are more resident throughout their whole life (Knutsen *et al*., 2011; André *et al*., 2016; Barth *et al*., 2019). Movement data from these tagged cod are not yet analysed, but may reveal more details about the spatiotemporal dynamics of the two ecotypes in inshore Skagerrak. Studies combining tagging efforts with population genetic analyses have the potential of resolving the functional roles that different ecotypes may play in the ecosystem, but such studies are still relatively rare (Henriksson *et al*., 2025).

#### 3.3.3. Population substructure within Eastern Baltic cod

The area around Åland, in the northern Baltic Sea, is one of very few areas in the Baltic Sea where large and mature cod in spawning condition are still found. In addition to them being substantially larger than cod in the Baltic Proper, otolith microchemistry suggests that Åland cod may have a local origin (Heimbrand & Limburg, 2025). To investigate potential genetic population substructure within EBC, we compared cod from Åland, as well as the even more northern sample from the Bothnian Sea, to the rest of the EBC samples. Pairwise *F*_ST_ estimates between all geographic samples, including these two northern Baltic samples, did not, however, indicate any genetic substructure within EBC, after FDR correction of the p-values. This was true both overall for collinear loci, as well as for the inversion genotypes (Table S5; Figure S2D). The genetic results, thus, suggest that EBC comprises a single, genetically admixed population, although this cannot be concluded without more in-depth genomic studies.

#### 3.3.4. Parentage assignment

In addition to population and inversion assignment, the SNP panel can also be used for parentage assignment. In a parallel study, we have used the same SNP panel to reliably infer parent-offspring pairs, as well as full- and half-sibling pairs (Henriksson *et al*., *in prep*.). This was done by calculating kinship coefficients as above, and assigning parentage with the COLONY software (v2.0.7.1; Jones & Wang, 2010). The 4000 loci in our SNP panel give sufficient power to detect kinship down to second degree family relations, following the recommendations of Manichaikul *et al*. (2010). By comparing against the genetic baseline samples presented herein, we were also able to assign parents and offspring to ecotype and estimate ecotype hybridisation rate (Henriksson *et al*., *in prep*.).

The panel is also being applied in a restocking project, aiming to restore the EBC population (“ReCod”). The project is rearing and releasing millions of cod larvae in the Baltic Sea, and by genotyping the broodstock cod together with wild-caught juvenile cod from this area, it is possible to assess whether the release of larvae enhances the recruitment of juveniles.

### 3.4. Considerations

#### 3.4.1. Loci for population assignment

The SNP panel was created by combining several smaller population-diagnostic panels (e.g., Weist *et al*., 2019; Henriksson *et al*., 2023), and these different subpanels can be used to answer specific questions or to make direct comparisons with previous studies. We still, however, recommend future users to apply the full panel of collinear loci (*i*.*e*., loci outside of inversions) for population assignment, as the clearest population assignments were obtained using all collinear loci. In addition, our inversion scans indicated patterns of LD in additional genomic regions potentially representing small chromosomal inversions (Figure S1; *cf*. Berg *et al*., 2015; Barth *et al*., 2017; Hoff *et al*., 2024). The population genetic clustering presented herein was not affected by the patterns of LD displayed in these regions, but caution is required if the SNP panel is used for more fine-scaled explorations of population structure.

#### 3.4.2. Small-scale studies

The case studies presented in Section 3.3 exemplify how research projects with small numbers of samples can benefit from comparisons against the genetic baseline dataset. As long as the same SNP panel is used, it is possible to combine the genotypic data from smaller studies together with the genotypes of the baseline dataset. By using the individuals in the genetic baseline as a reference for the three populations (offshore cod, coastal/WBC, and EBC), the discriminatory power and accuracy of population assignment in these smaller studies improves substantially. Platform effects are minimised when the same genotyping platform is used in multiple studies, but future users will still need to account for batch effects. While such batch effects cannot always be controlled explicitly, replicating individuals across multiple genotyping rounds (positive controls) can help estimate the magnitude of such effects.

#### 3.4.3. Limitations of the SNP panel

While we demonstrate that the SNP panel has a multitude of different applications, there are always limitations to a relatively small SNP panel such as this. As it is a high-graded panel, it is not an optimal tool for explorative population genomics. The SNP panel was developed as a diagnostic tool for assignment of known populations in the waters surrounding Sweden, meaning it may have limited power to infer more local (unknown) population structure on finer geographical scales. For such efforts, we recommend that whole-genome resequencing techniques are applied instead.

## 4. Conclusions

Overall, we demonstrate the applicability of a new SNP panel to assignment of population of origin in Atlantic cod from Swedish seas. The utilisation of fish captured in regular fisheries-independent surveys enables cost-efficient and standardised sample collection that can easily be replicated and applied for future genetic monitoring programs. Our results establish a genetic baseline for cod in the waters surrounding Sweden, but we also demonstrate the value and utility of this baseline outside of genetic monitoring. The baseline can be useful for research projects with small sample sizes, as well as for any study with research questions that require assignment of population-of-origin, inversion genotypes, or kinship. It also shows promise as a tool for wildlife forensics and individual recognition. In summary, the SNP panel and the genetic baseline presented herein enable standardising the identification and genetic monitoring of Atlantic cod populations in Sweden and surrounding marine regions.

## Acknowledgements

This project was funded by the Swedish Agency for Marine and Water Management through grants “1.2 Miljöövervakning med mera” (Dnr 1717-2022) and “1:11. Åtgärder för havs-och vattenmiljö” (Dnr 2066-2023). We thank Linda Laikre, Leif Andersson, and Mats Pettersson for their contributions to the conceptualisation and coordination of the project and the technical development of the multi-species SNP chip. We are also grateful to Olga Ortega-Martinez, Ulf Bergström, and Yvette Heimbrand for their help with selecting and providing samples.

## Statements & Declarations

### Author contributions

Conceptualisation: SH, CA, RP, and KJ. Data curation: SH and RP. Formal analysis: SH. Funding acquisition: KJ. Investigation: SH. Methodology: SH, RP, and CA. Project administration: KJ. Resources: CA and HW. Supervision: CA. Visualisation: SH. Writing – original draft: SH. Writing – review & editing: SH, CA, HW, and KJ.

### Data availability

Metadata for the cod samples and the SNP panel used to establish the genetic baseline are available as Tables S1 and S2, respectively. Genotype data and an R script for statistical analysis will be deposited on a data repository upon acceptance for publication.

