## Supplementary material for "Establishing a baseline for standardised genetic monitoring of Atlantic cod (*Gadus morhua*) in Sweden": Genetic_Monitoring_Cod_SupplementaryFigures.docx

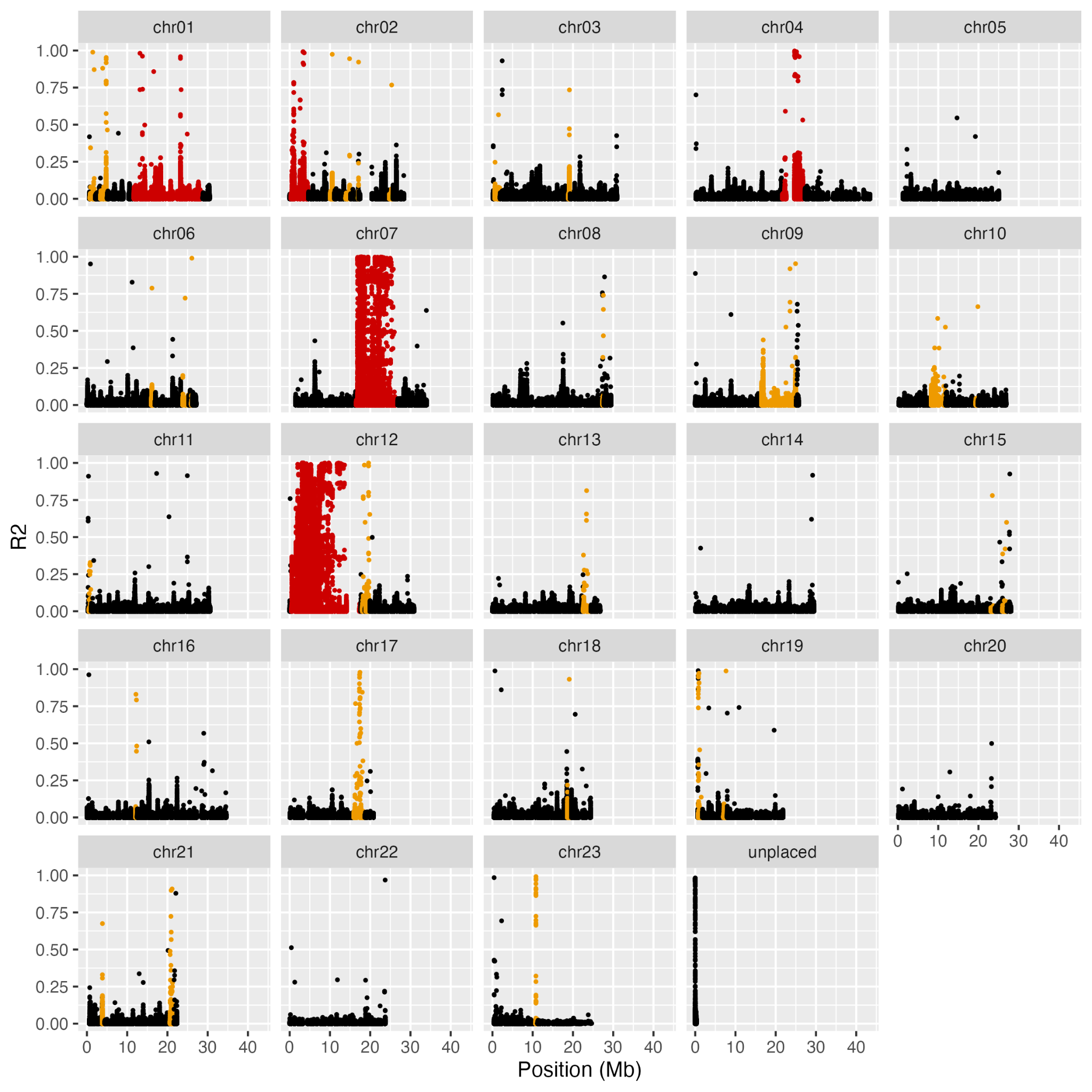

**Figure S1.** Linkage disequilibrium (LD) across the 23 chromosomes. The x axis shows the chromosomal position of all loci, the y axis shows the R^2^ value of all locus pairs, and colours indicate whether a locus is located within an inferred inversion (red), potential inversion (orange), or outside of any inversion (black). Note that the large inversion on chromosome 1 was not inferred in the present study, but by Aasegg Araya et al. (2025).

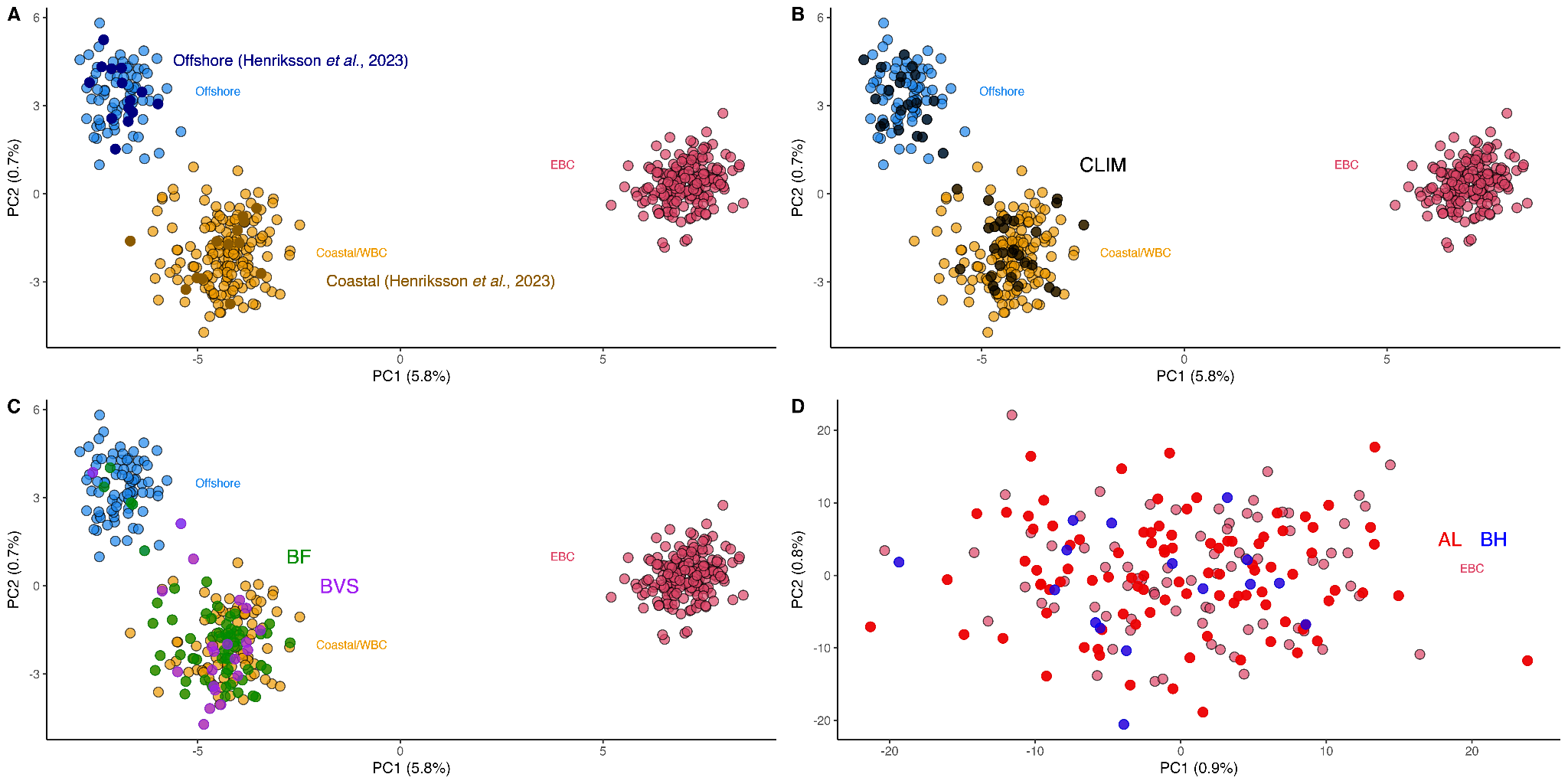

**Figure S2.** Principal component analyses (PCA) using collinear loci, with specific samples highlighted: **A)** previously ecotype-assigned cod from Henriksson et al. (2023), **B)** cod included in the climate stressor experiment by Perry et al. (2024; CLIM), **C)** large cod from inshore Skagerrak (BF = Byfjorden; BVS = Bovallstrand), and **D)** cod from Åland (AL) and the Bothnian Sea (BH) in the Eastern Baltic. Note that the PCAs in **A-C** include all samples, while the PCA in **D** was performed on Eastern Baltic cod (EBC) only.

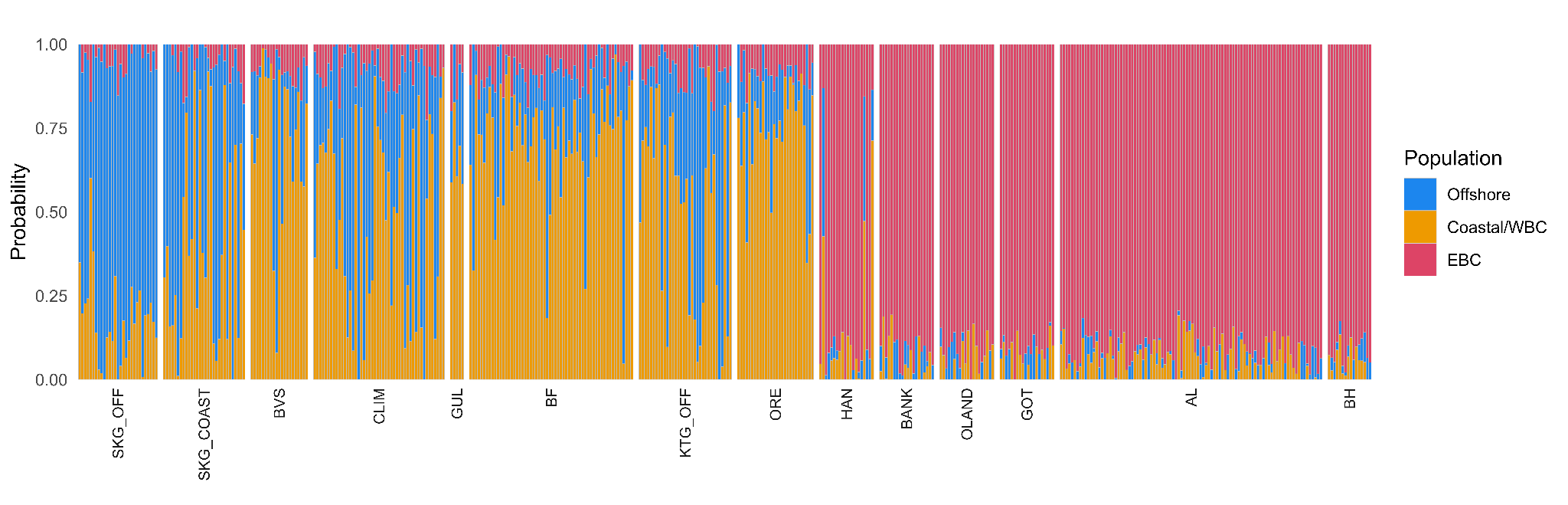

**Figure S3.** Ancestry coefficients for all individuals, calculated using sparse non-negative matrix factorisation (sNMF) at collinear loci. Samples are sorted from west (left) to east (right). Bar height represents the probability of an individual having a specific ancestry, with colour indicating the three ancestries.

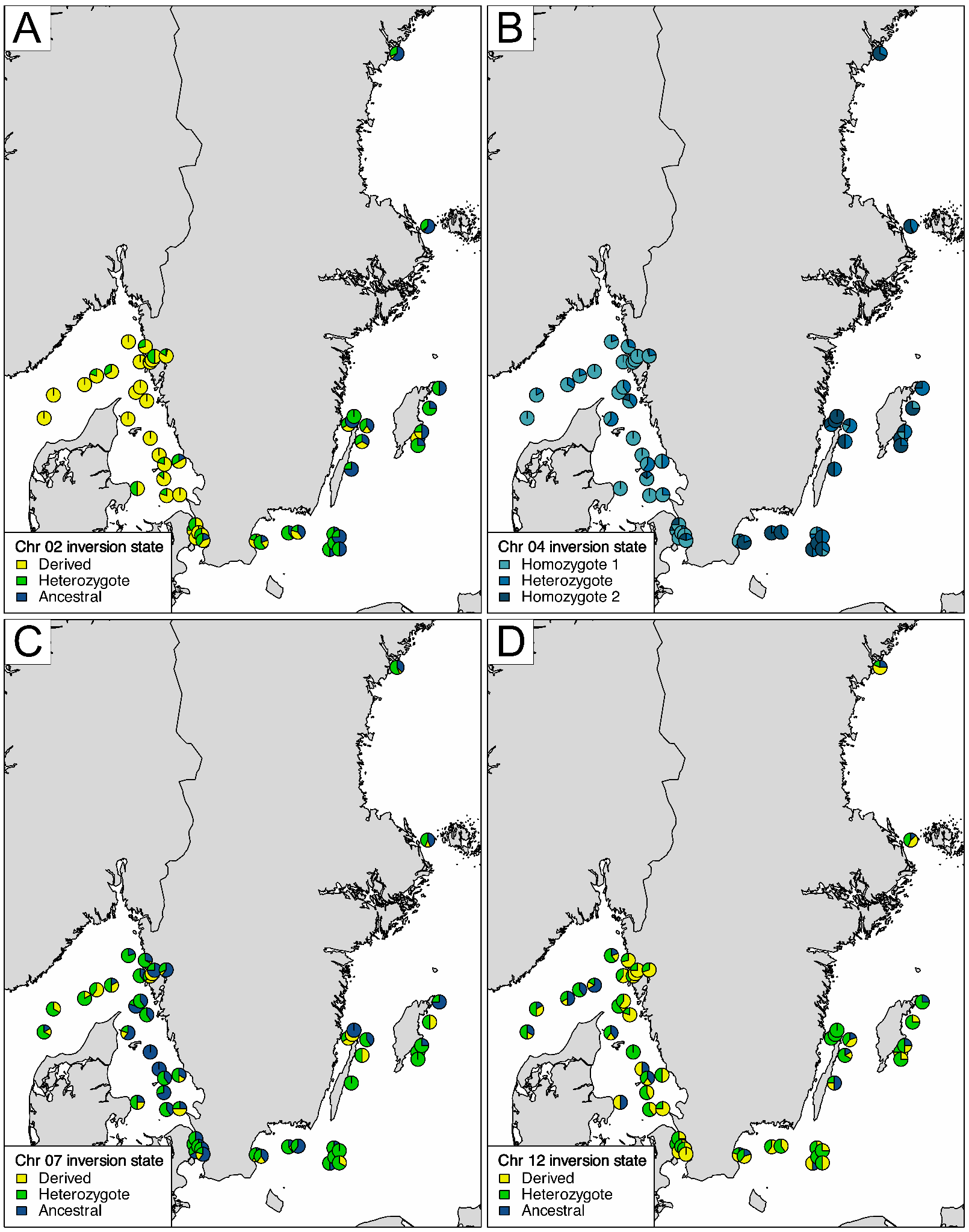

**Figure S4.** Inversion genotype frequencies across geographic locations for chromosomes **A)** 2, **B)** 4, **C)** 7, and **D)** 12. The relative size of the pie slices indicates the relative proportions of each population at each locality.

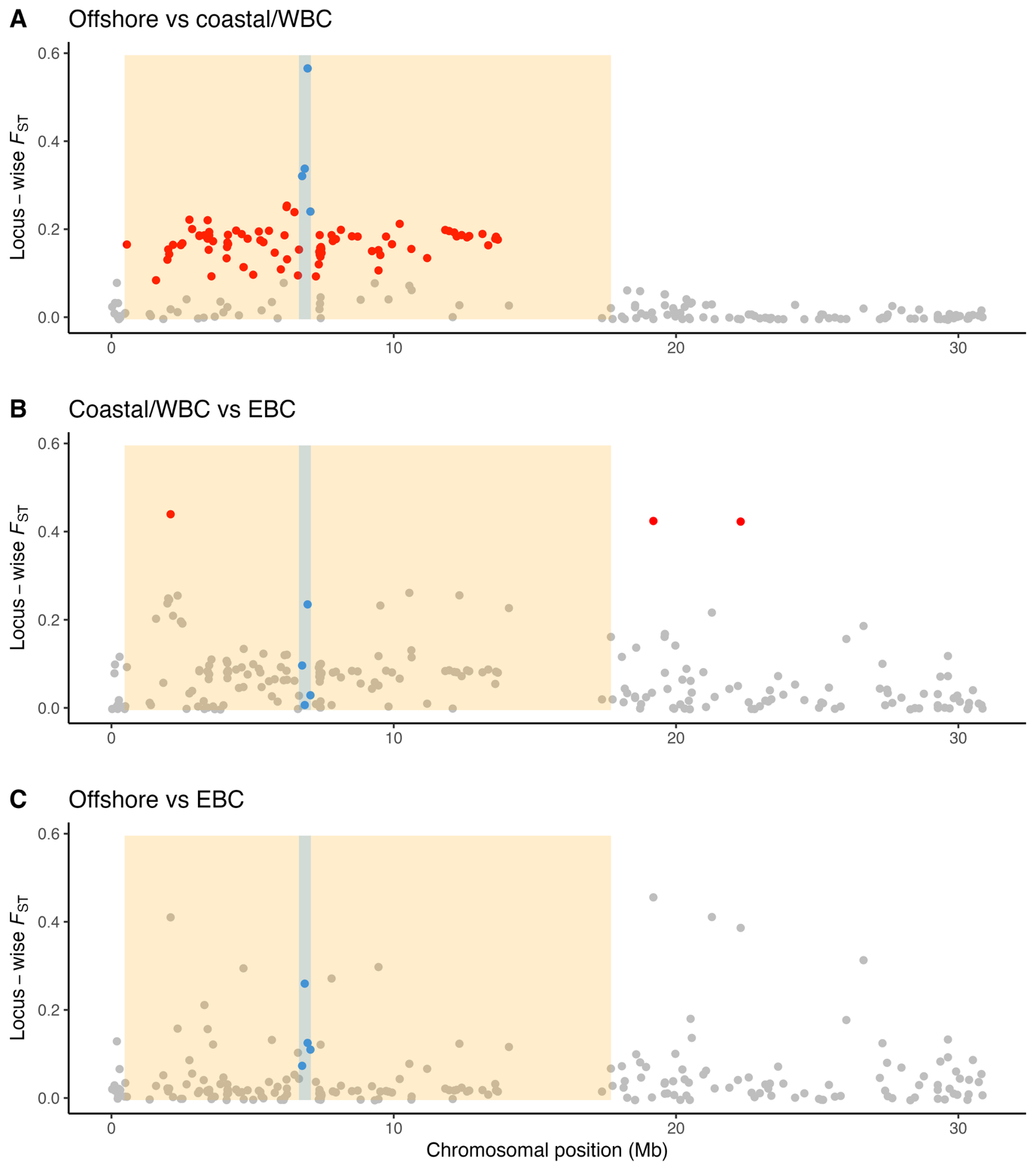

**Figure S5.** Pairwise *F*_ST_ across chromosome 12 for **A)** offshore vs coastal/Western Baltic cod (WBC), **B)** coastal/WBC vs Eastern Baltic cod (EBC), and **C)** offshore cod vs EBC. The region of the chromosomal inversion is highlighted in orange, and outlier loci are indicated with red points. The region of the potential double crossover, inferred by Matschiner et al. (2022), and the SNP loci located therein, are highlighted in blue.
